# Anterior hypothalamic nucleus drives distinct defensive responses through cell-type-specific activity

**DOI:** 10.1101/2024.09.25.615020

**Authors:** Cindy Yookyung Hong, Jessica Sofia Din, Hannah Chang, Jee Yoon Bang, Jun Chul Kim

## Abstract

Innate defensive behaviors, such as freezing, fleeing, and fighting, are essential for survival, enabling animals to effectively respond to predatory threats. These behaviors involve a complex interplay of sensory processing, decision-making, and motor output. As a core component of the medial hypothalamic defense system, the anterior hypothalamic nucleus (AHN) is a key brain region implicated in orchestrating innate defensive responses. Although the AHN is predominantly GABAergic, it also contains a smaller population of excitatory neurons, reflecting a sophisticated balance between inhibitory and excitatory signaling within this region. However, despite its importance, the specific behavioral functions of these diverse neuronal populations have not been systemically examined. In this study, we utilized fiber photometry and optogenetic stimulation to investigate the roles of AHN GABAergic, glutamatergic, and CaMKIIa+ neuronal activities in mediating innate defensive behaviors. Our results indicate that AHN GABAergic neurons mediate anxiety-associated investigatory behaviors, likely facilitating risk assessment during the pre-encounter stage. Conversely, AHN glutamatergic neurons drive escape initiation and freezing responses associated with the post-encounter stage. The AHN CaMKIIa+ neurons, which exhibit significant heterogeneity, suggest a more nuanced role, potentially balancing escape and freezing responses. By elucidating the functional specialization of different AHN neuron subtypes, this study provides a foundation for future investigations into the neural circuits underlying innate defensive behaviors and its dysregulation in neuropsychiatric conditions characterized by dysregulated responses to threats, such as PTSD and panic disorder.

## INTRODUCTION

When faced with threats, prey species first engage in a complex process of risk assessment behaviors, evaluating environmental factors such as distance to threat and the availability of safe shelters^1–3^. This assessment then informs the timely execution of defensive strategies—freezing, escape (flight), or defensive attack (fight) —tailored to the dynamically changing levels of threat^1,4^. Identifying the neural substrates underlying these behaviors is essential to understanding how animals dynamically transition and adapt between different defensive strategies, reflecting the complex interplay of perception, decision-making, and motor actions.

Three hypothalamic structures within the medial hypothalamic zone (MHZ) – the anterior hypothalamic nucleus (AHN), the dorsomedial part of the ventromedial hypothalamus (VMHdm), and the dorsal premammillary nucleus (PMd) – have been identified as critical drivers of innate defensive behaviors in response to natural predators^5–7^. The VMHdm and PMd, as glutamatergic excitatory structures, have well-established roles in mediating escape and freezing behaviors to both innate and conditioned threats^8–10^. In contrast, the AHN, with its predominantly GABAergic composition and a smaller population of glutamatergic neurons, presents a more complex picture regarding its role in defensive behaviors^11–14^. To fully understand the intricate role of the AHN in mediating defensive behaviors, a systematic characterization of the specific functions of its diverse neuronal populations is necessary. Therefore, our study aimed to investigate the roles of three molecularly distinct cell types within the AHN - GABAergic, glutamatergic, and heterogeneous CaMKIIa+ neurons - in mediating innate defensive behaviors.

While all three AHN neuronal populations – GABAergic, glutamatergic, and CaMKIIa+ – exhibited dynamic responses to a live predator, the timing of their activity varied across the cascade of approach and escape behaviors. AHN CaMKIIa+ neurons partially co-express GABA, suggesting potential functional heterogeneity within this population. Optogenetic stimulation experiments revealed that activation of AHN GABAergic neurons promotes thigmotaxis and increased sniffing during exploration, likely promoting risk assessment or investigatory behaviors. In contrast, activation of AHN glutamatergic and heterogeneous CaMKIIa+ neurons elicited flight or freezing responses. Lastly, we showed that activation of all three cell-types in the AHN is aversive.

Taken together, our study suggests a functional antagonism between two main cell types within the AHN. GABAergic AHN neurons promote anxiety-associated investigatory behaviors during the pre-encounter stage of threat assessment, whereas glutamatergic AHN neurons drive immediate escape initiation and freezing responses during the post-encounter stage, potentially associated with emotional states of panic and fear. Overall, we propose a unique role for the AHN within the MHZ, acting as a critical structure for regulating the intricate balance of innate defensive behaviors.

## RESULTS

### Predatory threat induces dynamic spatial response patterns in AHN cell types

The AHN is primarily composed of GABAergic neurons marked by vGAT or GAD2 expression^15^. However, it also contains a smaller population of excitatory glutamatergic neurons marked by vGLUT2 expression, as well as a subpopulation of neurons expressing the protein CaMKIIa^11,12,16,17^. To characterize the cell-type specific roles of the AHN in mediating innate defensive behaviors, we first investigated the activity patterns of distinct AHN neuronal populations during exposure to a live predator. We injected an adeno-associated virus (AAV) into the AHN to selectively express GCaMP6f, a genetically encoded calcium indicator, in AHN GABAergic neurons (AAV-hSyn-FLEX-GCaMP6f in VGAT-Cre mice), AHN glutamatergic neurons (AAV-hSyn-FLEX-GCaMP6f in VGLUT2-Cre mice) neurons, or AHN CaMKIIa+ neurons (AAV-CaMKIIa-GCaMP6f in C57Bl6 mice) (Figures 1A-1D and S1). To examine the response of these targeted cell populations to a live predatory threat, we performed fibre photometry during the predator-evoked avoidance task. In this task, mice were introduced to an open field arena containing a rat leashed to the corner of the apparatus (Figure 1E). During the approach phase, mice exhibited a combination of investigatory behaviors such as sniffing, reduced locomotion, and stretch-attend posture (SAP), indicating an initial risk assessment response (Figures S2A and S2B). This was then followed by a rapid escape away from the rat (Figure S2C). The GCaMP recordings during the behavioral paradigm were z-scored and analyzed to compare activity patterns across AHN cell types and assess the spatial dynamics of their responses.

**Figure 1.**
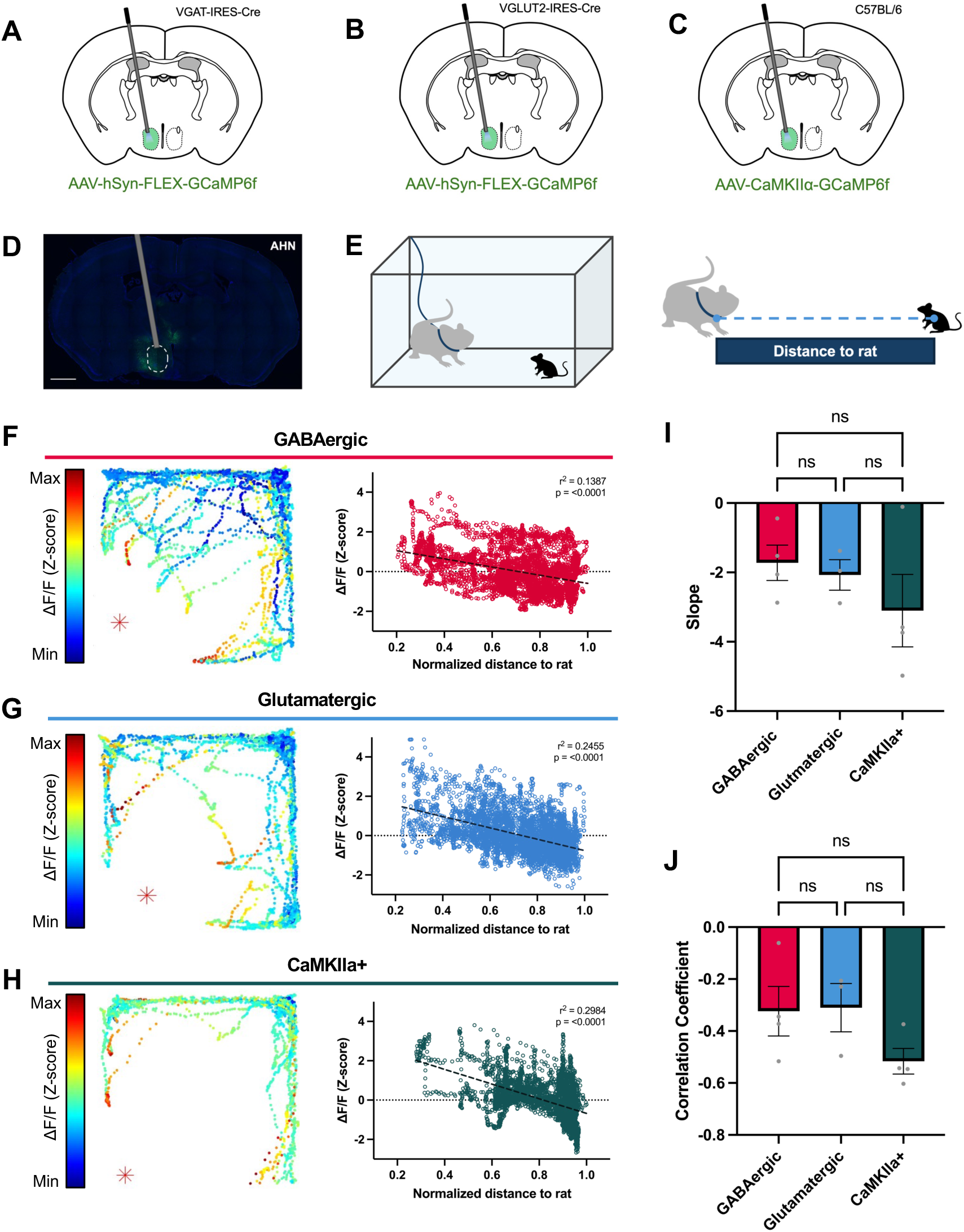
Predatory threat induces dynamic spatial response patterns in AHN cell types. (A-C) Schematic representations of fiber photometry recordings in (A) AHN GABAergic, (B) AHN glutamatergic, and (C) AHN CaMKIIa+ neurons (D) Representative image showing viral expression and optic fibre implantation in the AHN (E) Schematic representation of the predator-evoked avoidance task and spatial analysis during which mice were introduced to an open field arena with a rat leashed to the corner of the apparatus (F-H) Representative GCaMP6f trajectory heatmaps (left) depicting ΔF/F during the predator-evoked avoidance task, and correlational plot (right) showing the correlation between distance to threat and ΔF/F in (F) AHN GABAergic, (G) AHN glutamatergic, and (H) AHN CaMKIIa+ neuronal recordings. (I) Summary of slopes from correlational analyses between distance to threat and ΔF/F (AHN GABAergic n = 4, AHN glutamatergic n = 3, AHN CaMKIIa+ n = 4, One-way ANOVA F (2, 8) = 0.9295, P = 0.4336, with Tukey’s multiple comparisons test, p > 0.05). (J) Summary of correlation coefficients from correlational analyses between distance to threat and ΔF/F (AHN GABAergic n = 4, AHN glutamatergic n = 3, AHN CaMKIIa+ n = 4, One-way ANOVA F (2, 8) = 2.118, P = 0.1827, with Tukey’s multiple comparisons test, p > 0.05).

All three AHN neuronal populations – GABAergic, glutamatergic, and CaMKIIa+ – exhibited the highest GCaMP fluorescence levels closest to the rat, with negative correlations observed between distance to the threat and fluorescence intensity (Figures 1F-1H), indicating increased neural activity in response to the proximity of the threat. Notably, we found no significant differences in the slopes or correlation coefficients of these responses among the three neuron types (Figures 1I, 1J; n=4 mice/group for GABAergic and CaMKIIa+ neurons, n=3 mice for glutamatergic neurons). These results suggest that while all three cell types are similarly activated by predatory threats, their overall activity level might not be the sole determinant of risk assessment or escape initiation. Instead, the specific timing (i.e., temporal dynamics) and coordination of activity within these populations may be more important for orchestrating the behavioral sequence of defensive responses to predatory threats.

### AHN GABAergic, glutamatergic, and CaMKIIa+ neurons are activated at different stages of escape

To determine whether different AHN neuronal populations are preferentially activated at specific stages of escape response, we aligned GCaMP fluorescence signals relative to escape onset, defined as the moment mice began running away from the rat (Figures 2A and 2B). All three cell types displayed elevated GCaMP activity during approach and escape behaviors. However, the timing of this activity differed significantly. Both GABAergic neurons and CaMKIIa+ neurons displayed a peak in activity near escape onset, with above-baseline activity emerging before the initiation of escape. (Figures 2C, 2E, 2F). Notably, compared to CaMKIIa+ neurons, the activation pattern of AHN GABAergic neurons was broader, with sustained activity throughout approach and escape behaviors (Figures 2C, 2E, 2F). In contrast, AHN glutamatergic neurons exhibited a distinct peak in activity that occurred significantly later, after escape onset (Figures 2D and 2F). This suggests that AHN glutamatergic neurons may be preferentially activated during escape execution rather than the initial risk assessment phase.

**Figure 2.**
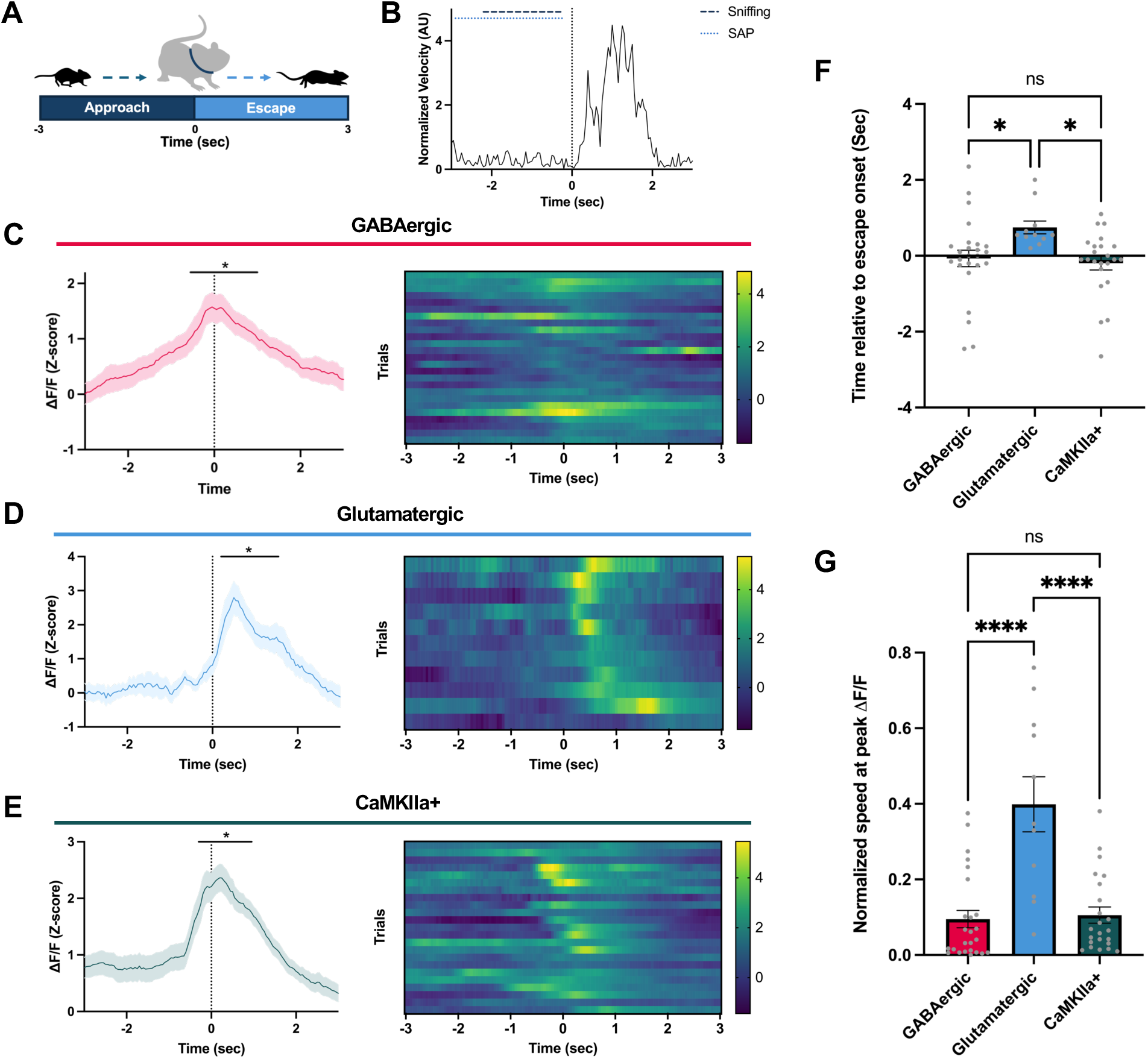
AHN GABAergic, glutamatergic, and CaMKIIa+ neurons are activated at different stages of escape. (A) Schematic representation of the temporal analysis of escape behavior in the predator-evoked avoidance task (B) Representative trace showing normalized velocity during approach and escape behavior. Sniffing and stretch attend posture (SAP) events are indicated by dashed and dotted lines, respectively (C-E) Average ΔF/F values during escape bouts aligned to escape onset (left) and heatmap of ΔF/F during individual escape bouts (right) for (C) AHN GABAergic (n = 25), (D) AHN glutmatergic (n = 11), and (E) AHN CaMKIIa-GCaMP6f (n = 23) neuronal recordings. The black line indicates timepoints where the change in ΔF/F from the start of the approach bout is statistically significant. (F) Time of peak ΔF/F value during escape bouts relative to escape onset (GABAergic n = 25, glutamatergic, n = 11, CaMKIIa+ = 23, One-way ANOVA F (2, 56) = 4.050, P = 0.0228, with Tukey’s multiple comparisons test, VGAT vs. VGLUT2 p < 0.05, VGAT vs. CaMKIIa p > 0.05, CaMKIIa vs. VGLUT2 p < 0.05) (G) Normalized speed at peak ΔF/F value during escape bouts (GABAergic n = 25, glutamatergic n = 11, CaMKIIa+ n = 23, One-way ANOVA F (2, 56) = 19.81, P < 0.0001, with Tukey’s multiple comparisons test, VGAT vs. VGLUT2 p < 0.0001, VGAT vs. CaMKIIa p > 0.05, CaMKIIa vs. VGLUT2 p < 0.0001)

Consistently, the running speed of mice at the peak GCaMP fluorescence was significantly higher for the AHN glutamatergic neurons, whereas no significant differences were observed between the AHN GABAergic and AHN CaMKIIa+ neurons (Figure 2G). Taken together, our results suggest unique roles for AHN GABAergic, glutamatergic, and CaMKIIa+ neurons in mediating innate defensive behaviors. AHN GABAergic activity arises during the risk assessment phase (i.e. approaching), while AHN glutamatergic neurons are preferentially activated later during the execution phase (i.e., running).

### Distinct GABAergic co-expression pattern in AHN CaMKIIa+ neurons

It is widely accepted that in cortical regions, CaMKIIa is expressed in glutamatergic cells^18,19^. However, recent findings suggest a unique anatomical and functional role of CaMKIIa+ neurons in hypothalamic regions^20,21^. To explore whether AHN CaMKIIa+ neurons exhibit distinct anatomical characteristics from GABAergic AHN neurons, we conducted immunohistochemistry staining on coronal brain sections expressing GFP in AHN GABAergic neurons (AAV-CAG-Flex-GFP in GAD2-IRES-Cre mice), AHN glutamatergic neurons (AAV-CAG-Flex-GFP in VGLUT2-IRES-Cre mice), and AHN CaMKIIa+ neurons (AAV-CaMKIIa-eYFP in C57BL/6 mice), using anti-GABA antibodies (Figure 3A).

**Figure 3.**
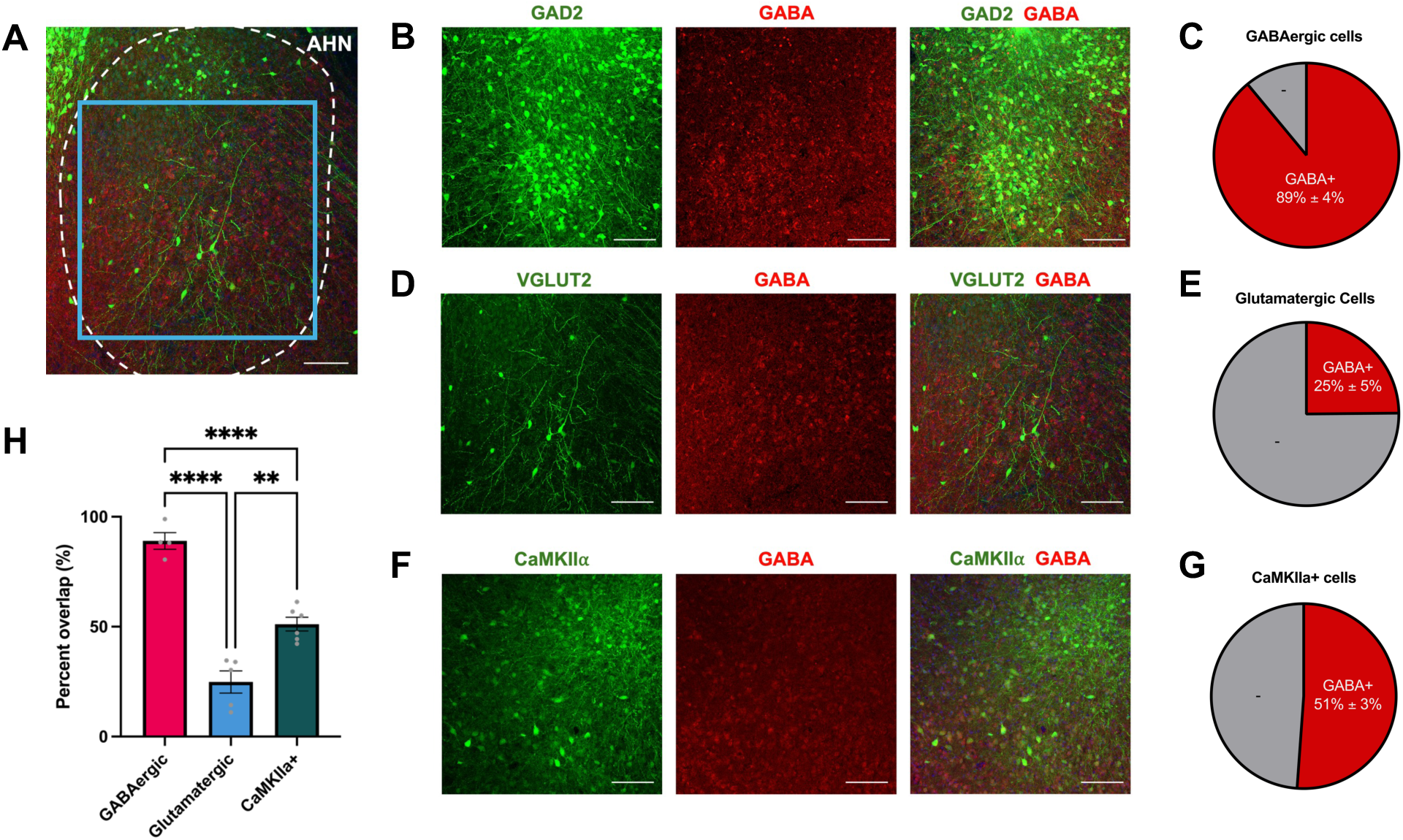
Distinct GABAergic co<u>-</u>expression pattern in AHN CaMKIIa+ neurons. (A) Representative micrograph showing viral expression of GFP in VGLUT2+ neurons and immunostaining for GABA (red) in the AHN (Scale bar, 100 um) (B) Representative micrographs for quantitative analyses showing AHN GAD2+:GFP (green) expression and GABA (red) immunostaining in the AHN (Scale bar, 100 µm) (C) Percentage of GABA+ and GABA- expression in AHN GAD2+ neurons (n = 2 mice) (D) Representative micrographs for quantitative analyses showing AHN VGLUT2+:GFP (green) expression and GABA (red) immunostaining in the AHN (Scale bar, 100 µm) (E) Percentage of GABA+ and GABA- expression in AHN VGLUT2+ neurons (n = 2 mice) (F) Representative micrographs for quantitative analyses showing AHN CaMKIIa+:eYFP (green) expression and GABA (red) immunostaining in the AHN (Scale bar, 100 µm) (G) Percentage of GABA+ and GABA- expression in AHN CaMKIIa+ neurons (n = 2 mice) (H) Percent overlap between GABA and AHN GAD2+, CaMKIIa+, and VGLUT2+ neurons (GAD2 n = 4 coronal slices, CaMKIIa n = 6 coronal slices, VGLUT2 n = 5 coronal slices, One-way ANOVA F (2, 12) = 56.49, P < 0.0001, with Tukey’s multiple comparisons test, GAD2 vs. CaMKIIa p < 0.0001, GAD2 vs. VGLUT2 p < 0.0001, CaMKIIa vs. VGLUT2 p < 0.01)

Our analysis revealed robust colocalization of GABA with AHN GABAergic neurons (Figures 3B and 3C), as expected. AHN glutamatergic neurons sparsely colocalized with GABAergic neurons (Figures 3D and 3E). Interestingly, we observed significant colocalization of GABA with AHN CaMKIIa+ neurons (Figures 3F and 3G), suggesting a potential heterogeneity within this population compared to the expected glutamatergic phenotype. Overall, the percent overlap between AHN GABAergic, glutamatergic, and CaMKIIa+ neurons with GABA immunoreactivity were significantly different between each cell-type, suggesting that these cell types exhibit distinct neurochemical and functional characteristics within the AHN (Figure 3H).

### Activation of AHN GABAergic, glutamatergic, and CaMKIIa+ neurons evoke distinct innate defensive behaviors

Both electrical and optogenetic stimulation of the entire pan-neuronal AHN have been demonstrated to elicit robust innate defensive behaviors in rodents, including escape running, jumping, and freezing behavior^9,13,22^. To test the cell-type specific role of AHN neurons in mediating innate defensive behaviors, we performed pan-neuronal or cell-type selective optogenetic stimulation targeting AHN GABAergic, glutamatergic, or CaMKIIa+ neurons. ChR2 or fluorophore control was virally expressed either throughout the entire AHN for pan-neuronal activation (AAV-hSyn-ChR2-eYFP in C57BL/6 mice), specifically in GABAergic neurons (AAV-ef1a-DIO-ChR2-eYFP in GAD2-IRES-Cre mice), glutamatergic neurons (AAV-ef1a-DIO-ChR2-eYFP in VGLUT2-IRES-Cre mice), or in CaMKIIa+ neurons (AAV-CaMKIIa-ChR2-mCherry in C57BL/6 mice) with optic fibres implanted bilaterally over the AHN (Figures 4A-D). Histological analyses confirmed the viral expression and verified the implantation of optic fibers in the AHN (Figures 4E and S3).

**Figure 4.**
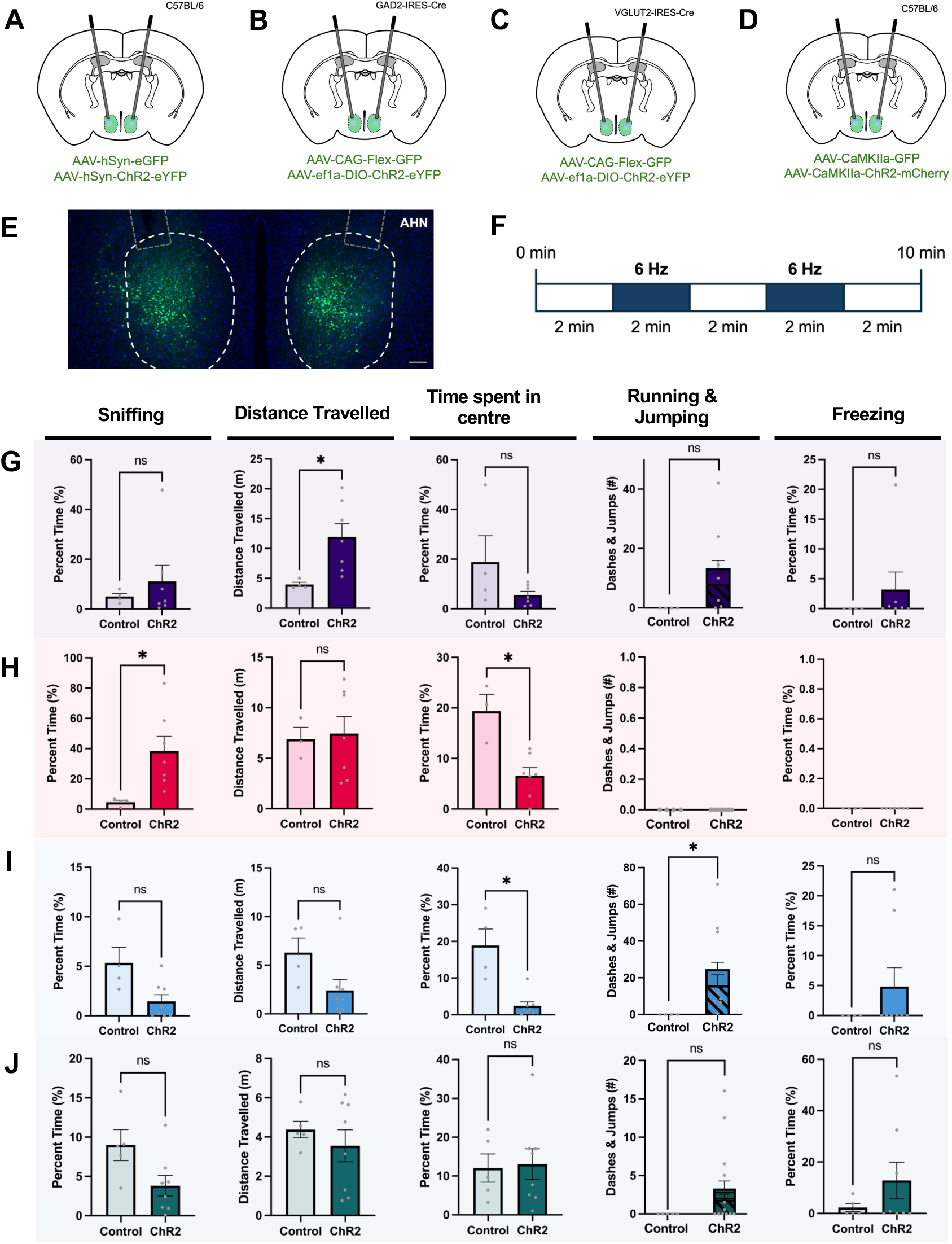
Activation of AHN GABAergic, glutamatergic, and CaMKIIa+ neurons evoke distinct innate defensive behavior. (A-D) Schematic representation of optogenetic stimulation in (A) AHN pan-neuornal, (B) AHN GABAergic, (C) AHN glutamatergic, and (D) AHN CaMKIIa+ neurons (E) Representative image of viral expression and optic fibre implantation in the AHN (Scale bar, 100 µm) (F) Schematic representation of optogenetic stimulation in an open field arena, with 6 Hz stimulations delivered at 2 minute light ON and OFF epochs (G) Comparisons of AHN pan-neuronal optogenetic stimulation effects between fluorophore controls (n = 4 mice) and ChR2 (n = 7 mice) mice on (from left to right) sniffing (Welch’s t-test p > 0.05), distance travelled (Welch’s t-test *p < 0.05), time spent in centre (Welch’s t-test p > 0.05), running and jumping (Welch’s t-test p > 0.05), and freezing (Welch’s t-test p > 0.05) responses during the light ON epoch. (H) Comparisons of AHN GABAergic neuronal optogenetic stimulation effects between fluorophore controls (n = 4 mice) and ChR2 (n = 7 mice) mice on (from left to right) sniffing (Welch’s t-test *p < 0.05), distance travelled (Welch’s t-test p > 0.05), time spent in centre (Welch’s t-test *p < 0.05), running and jumping, and freezing responses during the light ON epoch. (I) Comparisons of AHN glutamatergic neuronal optogenetic stimulation effects between fluorophore controls (n = 4 mice) and ChR2 (n = 8 mice) mice on (from left to right) sniffing (Welch’s t-test p > 0.05), distance travelled (Welch’s t-test p > 0.05), time spent in centre (Welch’s t-test *p < 0.05), running and jumping (Welch’s t-test *p < 0.05), and freezing (Welch’s t-test p > 0.05) responses during the light ON epoch. (J) Comparisons of AHN CaMKIIa+ neuronal optogenetic stimulation effects between fluorophore controls (n = 5 mice) and ChR2 (n = 8 mice) mice on (from left to right) sniffing (Welch’s t-test p > 0.05), distance travelled (Welch’s t-test p > 0.05), time spent in centre (Welch’s t-test p > 0.05), running and jumping (Welch’s t-test p > 0.05), and freezing (Welch’s t-test p > 0.05) responses during the light ON epoch.

**Figure 5.**
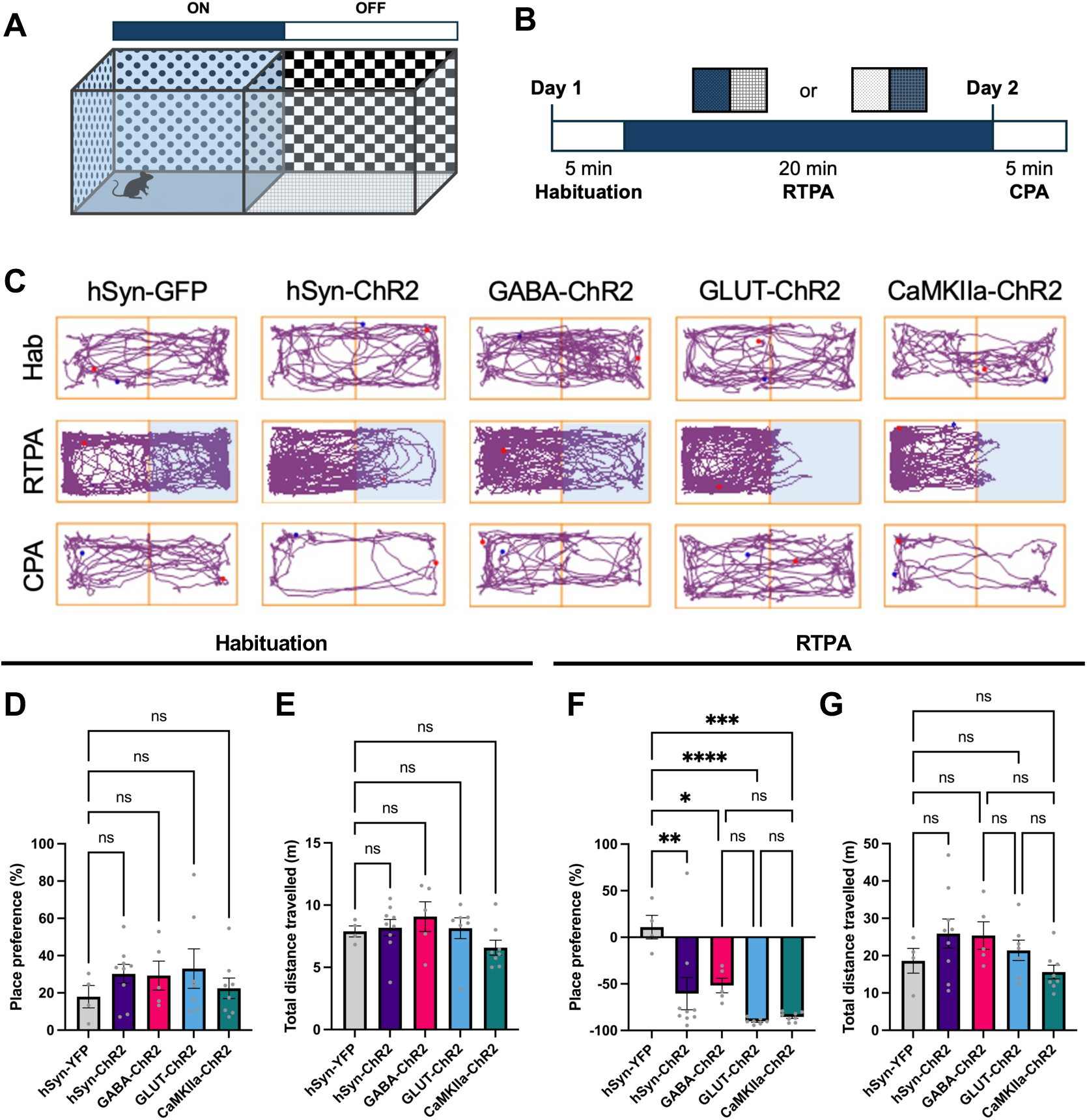
Activation of AHN GABAergic, glutamatergic, and CaMKIIa+ neurons is aversive. (A) Schematic diagram of the real-time place aversion apparatus. A two-chamber apparatus was used, each chamber with unique wall paper and floorings. (B) Schematic representation of the real-time place aversion (RTPA) and conditioned placed aversion (CPA) tests. Day 1 included an habituation session (5 min) followed by a RTPA test (20 min). Day 2 included a CPA test (5 min). (C) Representative trajectory plots for animals in each experimental cohort (left to right, hSyn-GFP, hSyn-ChR2, GABA-ChR2, GLUT-ChR2, and CaMKIIa-ChR2) during (top to bottom) habituation, RTPA, and CPA sessions. Light-coupled chambers are highlighted in blue. (D) Percent place preference for the preferred chambers during the habituation sessions by hSyn-GFP (n = 4 mice), hSyn-ChR2 (n = 9 mice), GABA-ChR2 (n = 5 mice), GLUT-ChR2 (n = 7 mice), and CaMKIIa-ChR2 (n = 8 mice) mice (One-way ANOVA F (4,28) = 0.6114, P > 0.05, with Tukey’s multiple comparisons test) (E) Total distance travelled during the habituation sessions by hSyn-GFP (n = 4 mice), hSyn-ChR2 (n = 9 mice), GABA-ChR2 (n = 5 mice), GLUT-ChR2 (n = 7 mice), and CaMKIIa-ChR2 (n = 8 mice) mice (One-way ANOVA F (4,28) = 1.374, P > 0.05, with Tukey’s multiple comparisons test) (F) Percent place preference for the stimulation chambers during the RTPA sessions by hSyn-GFP (n = 4 mice), hSyn-ChR2 (n = 9 mice), GABA-ChR2 (n = 5 mice), GLUT-ChR2 (n = 7 mice), and CaMKIIa-ChR2 (n = 8 mice) mice (One-way ANOVA F (4,28) = 8.830, P < 0.0001, with Tukey’s multiple comparisons test, *p < 0.05, **p < 0.01, ***p < 0.001, ****p < 0.0001) (G) Total distance travelled during the RTPA sessions by hSyn-GFP (n = 4 mice), hSyn- ChR2 (n = 9 mice), GABA-ChR2 (n = 5 mice), GLUT-ChR2 (n = 7 mice), and CaMKIIa-ChR2 (n = 8 mice) mice (One-way ANOVA F (4,28) = 1.976, P > 0.05, with Tukey’s multiple comparisons test)

Optogenetic stimulation was conducted in an open field arena at two-minute laser ON-OFF epochs for a total duration of ten minutes (Figure 4F). Two different stimulation frequencies (6 Hz or 20 Hz) were tested to assess potential frequency-dependent behavioral responses. Innate defensive behaviors such as sniffing, running, jumping, and freezing were manually scored and analyzed. Additional parameters including distance travelled and time spent in the center zone were also evaluated. Consistent with previous studies, we found that pan-neuronal stimulation of the entire AHN led to an increase in the total distance travelled and elicited a range of defensive behaviors, including running, jumping, and freezing, although not statistically significant (Figure 4G and S4A).

Given that the AHN is a predominantly GABAergic structure, we predicted that the observed effects of pan-neuronal activation of AHN, including running, jumping, and freezing could be reproduced by the stimulation of GABAergic neurons. Surprisingly, however, optogenetic stimulation of GABAergic neurons did not induce any running, jumping, or freezing (Figures 4H and S4B). Instead, GABAergic neuron activation, at both low and high frequencies, resulted in a striking increase in total sniffing time (Figures 4H and S2C), with animals exhibiting excessive ambulatory sniffing behavior (sniffing while moving) during stimulation (Figures S4E and S4F). Moreover, stimulation of AHN GABAergic neurons significantly decreased time spent in the center zone of the arena (Figures 4H and S4B), a behavior indicative of increase in anxiety, and increased locomotor activity along the walls of the apparatus (i.e., thigmotaxis) (Figure S4F). Thus, our findings suggest that GABAergic neurons within the AHN promote anxiety-associated investigatory behaviors involving sniffing and thigmotaxis. Additionally, the established role of AHN activity in escape running, jumping, and freezing responses may be mediated by other neuronal populations within the AHN, potentially including a small subpopulation of glutamatergic neurons and/or CaMKIIa+ neurons.

Consistent with our prediction, activation of AHN glutamatergic neurons evoked a significant increase in running and jumping behaviors, a significant reduction in total time spent in the center zone, and no significant changes in any other behavioral parameters (Figure 4I). Interestingly, high-frequency activation of these cell-types induced a significant increase in freezing behaviors and a significant decrease in sniffing behaviors (Figure S4C). These findings demonstrate that activation of AHN excitatory neurons robustly drives the execution of escape and freezing behaviors in a frequency-dependent manner.

Unlike AHN GABAergic and glutamatergic neurons, low frequency optogenetic stimulation of AHN CaMKIIa+ neurons did not trigger statistically significant changes in any behavioral parameters (Figure 4J). However, all stimulated mice in the experimental (ChR2) cohort (n=13) displayed visible increases in either running, jumping, or freezing behaviors (Figure 4J). High-frequency stimulation of CaMKIIa+ neurons resulted in a significant increase in both running and jumping behaviors (Figure S4D), suggesting a frequency-dependent modulation towards an escape response. This suggests activation of CaMKIIa+ neurons can evoke escape and freezing behaviors, but the lack of a clear dominant response might be due to the heterogeneity of these neurons. Taken together, our findings reveal distinct roles for different neuronal populations within the AHN in mediating innate defensive behaviors. Activation of GABAergic neurons promotes anxiety-associated investigatory behaviors, while activation of glutamatergic neurons drives escape initiation and execution.

### Activation of AHN GABAergic, glutamatergic, and CaMKIIa+ neurons is aversive

To probe the emotional valence associated with AHN GABAergic, AHN glutamatergic, and AHN CaMKIIa+ activation, we performed a real-time place aversion (RTPA) assay. Mice were first habituated to a two-chamber apparatus with distinct wallpaper and flooring in each chamber (Figure 6A). Following habituation, chamber preference was assessed, and the chamber where the mouse spent more time was designated as the stimulation chamber. During a subsequent 20-minute RTPA session, low-frequency stimulations were administered when mice entered the stimulation chamber. 24 hours after testing, mice were reintroduced into the same apparatus to assess conditioned place aversion (CPA) (Figure 6B).

During habituation, no significant differences in chamber preference and distance travelled were observed between the GFP control group and all four experimental cohorts (Figures 6C-6E). However, during the RTPA session, pan-neuronal and selective stimulations of the AHN, AHN GABAergic, AHN glutamatergic, and AHN CaMKIIa+ neurons evoked significant aversion away from the stimulation chamber with no changes in the total distance travelled (Figures 6F and 6G). While not statistically significant, activation of the AHN appeared to induce a trend towards mild conditioned place aversion of the stimulation chamber, with no changes in locomotor activity (Figures S5A and S5B). Taken together, our results demonstrate that stimulation of the AHN GABAergic, AHN glutamatergic, and AHN CaMKIIa+ neurons is aversive.

## DISCUSSION

From an ethological viewpoint, innate defensive behaviors vary across a predatory imminence continuum based on the dynamically changing interactions with predatory threat^23^. This continuum highlights three stages of predator encounters - pre-encounter, post-encounter and circa-strike – each corresponding to emotional states of anxiety, fear, and panic, respectively^23–26^. During the pre-encounter stage, prey animals engage in risk assessment behaviors to evaluate their surroundings and potential threats. Upon predator detection (post-encounter), they execute appropriate defensive strategies, including freezing, fleeing, or fighting, depending on the perceived threat level, threat proximity, and environment. Our study sheds light on the cell-type specific roles of the anterior hypothalamic nucleus (AHN) in orchestrating these behaviors.

Fiber photometry experiments revealed that while AHN GABAergic, glutamatergic, and CaMKIIa+ neurons all respond dynamically to predatory threat, the temporal activity pattern of each cell type is unique to different stages of approach and escape behaviors. Specifically, AHN GABAergic and CaMKIIa+ neurons are pre-emptively activated as mice approach a predatory threat prior to escape onset, whereas AHN glutamatergic neurons are activated almost exclusively during escape execution. These findings suggest that a cascade of cell-type specific activity within the AHN may orchestrate the full sequence of innate defensive behaviors, from initial threat assessment to the execution of defensive actions. It remains unclear, however, if these cell types respond exclusively to predatory threats or to other environmental cues associated with threat and safety. Recent studies by Laing et al. showed that AHN neurons indeed respond to contextual information associated with a live predator and to auditory cues associated with electrical foot shock during fear conditioning^12,27^. This suggests that AHN neurons can encode a broader spectrum of threat-related stimuli beyond immediate predator encounters, highlighting the need for further research to elucidate the full spectrum of stimuli that activate these AHN neuron populations and to determine their specific contributions to defensive behaviors in varied contexts.

Direct electrical and optogenetic stimulation of the entire AHN have been found to elicit a robust escape jumping and running^9,22^. However, recent studies targeting specific cell types reported highly mixed results, ranging from defensive attack, avoidance of novel stimuli, and escape initiation, challenging the previously held view of the AHN as a homogenous structure involved in promoting escape responses. Our cell type-specific optogenetic stimulation experiments revealed a distinct functional division within the AHN. The activation of GABAergic neurons within the AHN evoked anxiety-associated investigatory behaviors, likely serving as a counterbalance to the immediate escape drive during threat detection and risk assessment stage. GABAergic AHN neurons have recently been identified as drivers of defensive attack behaviors against predators^14^. Interestingly, however, we did not observe any defensive attack behaviors upon stimulation of GABAergic AHN neurons in the presence of a live rat (data not shown). This finding suggests that previously reported defensive attack behaviors may be an indirect consequence of excessive investigatory responses rather than a direct effect of GABAergic neuron activation.

In contrast to the risk assessment behaviors evoked by AHN GABAergic neurons, optogenetic stimulation of glutamatergic neurons in the AHN triggered escape running, jumping, and freezing behaviors, reflecting more immediate defensive strategies. Notably, varying frequencies of glutamatergic neuronal activation evoked distinct responses: low frequency stimulation evoked running and jumping whereas high frequency stimulation evoked freezing behaviors. This frequency-dependent modulation aligns with the concept of scalable emotional states underlying the control of defensive behaviors^28,29^. Consistent with our findings, a recent study has provided additional insight into the role of a specific population of parvalbumin (PV)-expressing glutamatergic neurons within the AHN in promoting escape behavior^12^. While stimulation of AHN PV neurons did not trigger immediate escape running or jumping, it induced avoidance behavior and increased locomotion speed during real-time place preference tests. Furthermore, ablation of AHN PV neurons resulted in delayed escape responses during a predator-looming task^12^. These findings suggest that AHN PV neurons likely represent a small subset of vGLUT2+ excitatory neurons and underscore the complexity of the AHN’s excitatory neuron population and their diverse roles in orchestrating defensive behaviors.

Growing evidence suggests that CaMKIIa+ neurons may represent an anatomically and functionally unique subpopulation in hypothalamic regions^20,21^. We found that a significant proportion (51%) of AHN CaMKIIa+ neurons are GABAergic. Consistently, unlike the distinct and robust responses evoked by AHN GABAergic and glutamatergic neurons, optogenetic stimulation of AHN CaMKIIa+ neurons did not elicit a clear dominant behavioral effect. This presents an intriguing possibility into the role of AHN CaMKIIa+ neurons in balancing intricate defensive behaviors, potentially modulating both inhibitory and excitatory responses depending on the context of predatory threat. To fully understand how AHN CaMKIIa+ neurons contribute to innate defensive behaviors, further investigation into their specific anatomical and physiological properties is warranted. This includes elucidating the distinct pathways through which these neurons exert their effects and determining how their activity integrates with other neural circuits involved in defensive responses.

Animals that rely heavily on smell (olfactory-oriented animals) engage in a specific behavior called odor-scanning during threatening situations as a form of risk assessment^30,31^. This involves actively sniffing and moving around their surroundings to gather information about potential predators and assess shelter availability, shelter location, and the likelihood of escape success^11,23^. Upon detecting a predator, prey animals weigh the above variables carefully to determine the most adaptive course of action. If there is no safe shelter nearby, they actively choose freezing over escape flight to avoid being detected by predators. However, if they have previously encountered a nearby shelter and remember its location, the defense strategy quickly switches from freezing to escape running toward the remembered shelter location^32^.

Overall, we demonstrate that activation of AHN GABAergic, glutamatergic, and CaMKIIa+ neurons evoke distinct behavioral responses, highlighting a functional division of labor within the AHN. AHN GABAergic neuronal activation increased thigmotaxis and ambulatory sniffing behaviors, indicative of anxiety-associated investigatory responses observed during the pre-encounter stage of threat assessment. In contrast, selective activation of AHN glutamatergic neurons evoked robust escape and freezing behaviors, reflecting immediate defensive responses characteristic of the post-encounter stage.

## ACKNOWLEDGEMENTS

We thank members of the Kim Laboratory, Ashleigh Brink, Dr. Kaori Takehara, Dr. Rutsuko Ito, and Dr. Laura Corbit for their valuable discussions regarding this work. This work was supported by operating grants to J.C.K. from the Canadian Institutes of Health Research (CIHR) (MOP 496401) and the Natural Sciences and Engineering Council of Canada (NSERC) (MOP 491009)

## AUTHOR CONTRIBUTIONS

Conceptualization, C.Y.H and J.C.K.: Methodology, C.Y.H and J.C.K.: Software, C.Y.H. and J.C.K.: Validation, C.Y.H. and J.C.K.: Formal Analysis, C.Y.H. and J.S.D.: Investigation, C.Y.H., J.S.D., and H.C.: Resources, C.Y.H., J.Y.B., and J.C.K.: Data Curation, C.Y.H.: Writing – Original Draft, C.Y.H. and J.C.K.: Writing – Review & Editing, C.Y.H. and J.C.K.: Visualization, C.Y.H.: Supervision, J.C.K.: Project Administration, C.Y.H. and J.C.K.: Funding Acquisition, J.C.K.

## METHODS

### Animals

Male and female C57BL/6 (Charles River), Gad2-IRES-Cre (Strain #010802; The Jackson Laboratory), Vgat-ires-Cre knock-in (C57BL/6J) (Strain #: 028862; The Jackson Laboratory), and Vglut2-ires-Cre knock-in (C57BL/6J) (Strain #: 028863; The Jackson Laboratory) mice at 12-25 weeks of age were used for behavioral experiments. All mice were single-housed in a 12-hour light/dark cycle with food and water provided ad libitum. Adult male SHR/NHsd rats (Charles River) at 23-50 weeks of age were used as live predators for the predator-evoked avoidance tests. All rats were single-housed and maintained in a 12-hour light/dark cycle with food and water provided ad libitum. Behavioral tests were performed during the light cycle. All experimental procedures were in accordance with the guidelines of the Canadian Council on Animal Care and the Local Animal Care Committee at the University of Toronto.

### Viral vectors and stereotaxic surgery

For all surgical procedures, animals were induced with 4% isoflurane and maintained with 2% isoflurane at an oxygen flow rate of 1 L/min. Ophthalmic ointments were applied to animals’ eyes throughout all surgeries. Meloxicam was provided for analgesic treatment before surgery and during post-operative recovery. The anterior hypothalamic area (AP: −0.85 mm, ML: −0.45 mm, DV: −5.2 mm) was infused with an adeno-associated virus (92 nl for fiber photometry experiments; 2 x 25.2 nl for optogenetic experiments) using a pulled glass needle and Nanoject II (Drummond Scientific) at a 46 nl/s rate.

Fiber optic cannulas (1.25 mm ferrule size, 200 µm core diameter, 0.37 NA, Neurophotometrics, RWD; 200 um core diameter, 0.39 NA, Thorlabs) were implanted in the AHN (AP: −0.85 mm, ML: −1.38 mm, DV: −5.1 mm, 10° towards the midline). All animals were handled for a minimum of 5 minutes for 3 days prior to behavioral testing.

The following viruses were used:

- AAV8-hSyn-GCaMP6f
- AAV9-CaMKIIa-GCaMP6f
- AAV9-hSyn-FLEX-GCaMP6f
- AAV8-hSyn-eGFP
- AAV9-CAG-FLEX-GFP
- AAV5-CaMKIIa-eYFP
- AAV9-hSyn-ChR2-eYFP
- AAV5-CaMKIIa-ChR2-mCherry
- AAV9-ef1a-DIO-ChR2-eYFP

### Fiber photometry recording and data analysis

A fiber photometry system (FP3002; Neurophotometrics) was used to record GCaMP6f signals in AHN neurons. Two LED beams were delivered at excitation wavelengths of 415 nm and 470 nm and interleaved at 40 Hz to record isosbestic and calcium-dependent activities, respectively. The light was passed through a single silica patch cord (200 µm core diameter, 0.37 NA, 3 m long; Thorlabs) which was connected to the implanted fiber optic cannula. LED intensity was measured at 50 µW using a power meter (PM100USB; Thorlabs) prior to all photometry experiments. Photometry data were acquired using a custom Bonsai script and analyzed in MATLAB. To account for calcium-independent signal fluctuations (415 nm), a linear interpolation model was fitted to the 20 Hz recorded data, approximating LED signal decay and fluorophore bleaching. Subsequently, calcium-dependent data (470 nm) were normalized by dividing them by the scaled model, correcting for bleaching. Finally, the entire dataset was z-scored to standardize measurements.

### Predator-evoked avoidance

Animals were introduced to an opaque plexiglass chamber (40 cm x 40 cm x 40 cm) with a rat leashed to one corner of the apparatus. Mice were left to freely explore the arena for a total of five minutes.

### Optogenetic manipulation

Blue light (473 nm, 6 Hz or 20 Hz) was produced using a diode-pumped solid-state laser (Laserglow). A patch cable (200 µm core diameter, 0.37 NA; Doric Lenses) was connected to a 1 x 2 optical communicator (Doric Lenses) to divide the light path from the laser into two optical fibers attached to the implanted optic fibers. Light power was measured at a power intensity of 5 mW at the optic fiber tip using a power meter (PM121D; Thorlabs). Light delivery was controlled through either a custom Arduino circuit paired with Bonsai programming for open field manipulations or ANY-MAZE (Stoelting Co.) for RTPA tests.

### Optostimulation of AHN neurons in open field

Mice were habituated to the tethering cable for fifteen minutes in their home cage and then introduced to a clear plexiglass chamber (50 cm x 50 cm x 20 cm), where low and high frequency optostimulation effects were compared. Animals were habituated to the apparatus for two minutes. Subsequently, two 2-minute photostimulations (5 mW, 6Hz, 5 ms pulse width) were delivered at two-minute laser ON-OFF intervals for a total of ten minutes. One week after testing, the behavioral test was repeated with 20 Hz stimulations. ANY-MAZE (Stoelting Co.) was used to determine the distance travelled and time spent in center, in addition to blindly and manually scoring sniffing, rearing, grooming, running, jumping, and freezing behaviors.

### Real-time place aversion (RTPA) and conditioned place aversion (CPA)

A custom-made two-chamber apparatus (45 cm x 20 cm x 35 cm) with distinct visual contexts and textured floorings was used. Animals were habituated to the tethering cable for 15 minutes in their home cages before being introduced to the apparatus. Mice were habituated to the apparatus for 5 minutes, and the place preference in each chamber was assessed. The preferred chamber was selected as the stimulation chamber. Animals received 6 Hz stimulation upon entering the stimulation chamber during a 20-minute RTPA test. After 24 hours, animals were returned to the two-chamber apparatus, and the place preference during the first five minutes was assessed as a measure of retrieval CPA memory. ANY-MAZE (Stoelting Co.) was used to assess total time spent in each chamber, distance travelled, and track plots. The place preference index during the apparatus habituation was calculated using the following equation: PPI (%) = [(Time spent in preferred chamber – time spent in less preferred chamber)/Total time spent in both chambers] x 100. Conditioned place aversion during the CPA sessions was calculated by subtracting the PPI during the apparatus habituation from the PPI during the CPA session.

### Histology

Mice were anesthetized with Avertin and transcardially perfused with 0.1 M phosphate-buffered saline (PBS, pH7.4), followed by 4% paraformaldehyde (PFA). The brain tissues were removed and fixed in 4% PFA for 24 hours and immersed in 30% sucrose solution for 48 hours. Free-floating coronal sections (40 µm) were prepared with a cryostat (Leica, Germany). For confirmation of cell-type-specific viral labelling, the sections were washed in PBS, permeabilized with PBS containing 0.3% Triton X-100 (PBS-T), and blocked with 5% normal donkey serum (Jackson ImmunoResearch). The tissue sections were incubated with primary antibody (rabbit anti-GABA, 1:1000 in PBS-T, Sigma Aldrich) at 4 °C for 48 hours. The sections were rinsed with PBS-T and incubated with secondary antibody (Alexa Fluor 594-conjugated donkey anti-rabbit, 1:500, Jackson ImmunoResearch) at room temperature for 2 hours. The sections were rinsed with PBS, mounted onto slides, stained with DAPI solution (1 ug/ml in PBS), and cover-slipped with Aquamount (Polysciences, Inc., Warrington, PA). For fiber photometry and optogenetics experiments, the sections were washed in PBS, mounted onto slides, stained with DAPI solution (1 ug/ml in PBS), and cover-slipped with Aquamount (Polysciences, Inc., Warrington, PA) without additional staining.

For viral expression analyses and fibre implantation target confirmation, images were captured using a Zeiss upright fluorescent microscope with a 20 x objective. All cell-counting images were captured using a Leica TCS SP8 confocal microscope. Z-stack images were captured using a 20 x oil-immersion objective with a HyD detector for GFP (500 – 540 nm emission, 100% gain, 1.0 AU pinhole diameter) and RFP (590 – 630 nm emission, 100% gain, 1.0AU pinhole diameter), and a photomultiplier tube detector for DAPI (430 – 475 emission, 800V gain, 1.0 AU pinhole diameter). Images were processed using Fiji.

### Statistical analysis

All statistical analyses were performed using GraphPad Prism 10 (GraphPad Software). For time comparisons during the rat-evoked conditioned place aversion paradigm, the significance of the change in PPI and time spent was evaluated using a paired t-test. For photometry experiments, a one-way ANOVA was performed to compare GCaMP signals across the three chambers. To evaluate statistical significance of behaviors in the open field optogenetic stimulation experiment, a two-way repeated-measures ANOVA (2-way RM ANOVA) followed by Sidak’s multiple comparison test was performed to compare the interaction between treatment groups (GFP vs ChR2) and stimulation frequencies (6 Hz vs 20 Hz). A one-way ANOVA (1-way ANOVA) followed by a Tukey’s multiple comparison test was used for RTPA analysis between different treatment groups and cell types (hSyn-GFP, hSyn-ChR2, GAD2-ChR2, CaMKIIa-ChR2, VGLUT2-ChR2). Significance markers were defined as the following: * p<0.05, ** p<0.01, *** p<0.001, ****p<0.0001.

## SUPPLEMENTAL INFORMATION

Document S1. Figures S1-S5

**Figure S1. Viral expression and locations of fibre optic cannulae in the AHN for fibre photometry recordings**

(A) Viral expression depicted by green blots on coronal sections of the mouse brain atlas. Fibre optic cannulae implantation sites indicated by circles.

**Figure S2. AHN GABAergic, glutamatergic, and CaMKIIa+ neurons are activated during innate defensive behaviors**

(A) Percent animals demonstrating stretch attend postures (SAP) during approach and escape bouts (n = 21)

(B) Percent animals demonstrating sniffing behaviors during approach and escape bouts (n = 21)

(C) Normalized velocity during approach and escape bouts (n = 36)

**Figure S3. Viral expression and locations of fibre optic cannulae in the AHN for optogenetic studies**

(A) Viral expression depicted by green blots on coronal sections of the mouse brain atlas. Fibre optic cannulae implantation sites indicated by circles.

**Figure S4. High frequency activation of AHN GABAergic, glutamatergic and CaMKIIa+ neurons evoke distinct innate defensive behaviours**

(A) Comparisons of AHN pan-neuronal optogenetic stimulation effects between fluorophore controls (n = 4 mice) and ChR2 (n = 7 mice) mice on (from left to right) sniffing (Welch’s t-test p > 0.05), distance travelled (Welch’s t-test p > 0.05), time spent in centre (Welch’s t-test *p < 0.05), running and jumping (Welch’s t-test p > 0.05), and freezing (Welch’s t-test p > 0.05) responses during the light ON epoch.

(B) Comparisons of AHN GABAergic neuronal optogenetic stimulation effects between fluorophore controls (n = 4 mice) and ChR2 (n = 7 mice) mice on (from left to right) sniffing (Welch’s t-test *p < 0.01), distance travelled (Welch’s t-test p > 0.05), time spent in centre (Welch’s t-test *p < 0.05), running and jumping, and freezing (Welch’s t-test p > 0.05) responses during the light ON epoch.

(C) Comparisons of AHN glutamatergic neuronal optogenetic stimulation effects between fluorophore controls (n = 4 mice) and ChR2 (n = 8 mice) mice on (from left to right) sniffing (Welch’s t-test *p < 0.05), distance travelled (Welch’s t-test p > 0.05), time spent in centre (Welch’s t-test p > 0.05), running and jumping (Welch’s t-test p > 0.05), and freezing (Welch’s t-test **p < 0.01) responses during the light ON epoch.

(D) Comparisons of AHN CaMKIIa+ neuronal optogenetic stimulation effects between fluorophore controls (n = 5 mice) and ChR2 (n = 13 mice) mice on (from left to right) sniffing (Welch’s t-test p > 0.05), distance travelled (Welch’s t-test p > 0.05), time spent in centre (Welch’s t-test p > 0.05), running and jumping (Welch’s t-test **p < 0.01), and freezing (Welch’s t-test p > 0.05) responses during the light ON epoch.

(E) Representative image of a mouse in sniffing posture during GABAergic-ChR2 optogenetic simulation

(F) Representative top-down trajectory plots during a light ON stimulation session (20 Hz) for GABAergic-GFP and GABAergic-ChR2 mice

**Figure S5. Activation of AHN GABAergic, glutamatergic, and CaMKIIa+ neurons induces mild conditioned place aversion**

(A) Percent place preference for the stimulation chambers during the CPA sessions by hSyn-GFP (n = 4 mice), hSyn-ChR2 (n = 9 mice), GABA-ChR2 (n = 5 mice), GLUT-ChR2 (n = 7 mice), and CaMKIIa-ChR2 (n = 8 mice) mice (One-way ANOVA F (4, 28) = 2.009, P > 0.05, with Tukey’s multiple comparisons test, p > 0.05)

(B) Total distance travelled during the CPA sessions by hSyn-GFP (n = 4 mice), hSyn-ChR2 (n = 8 mice), GABA-ChR2 (n = 5 mice), GLUT-ChR2 (n = 7 mice), and CaMKIIa-ChR2 (n = 8 mice) mice (One-way ANOVA F (4, 27) = 1.328, P > 0.05, with Tukey’s multiple comparisons test, p > 0.05)

