## Supplemental Figures for "Anterior hypothalamic nucleus drives distinct defensive responses through cell-type-specific activity"

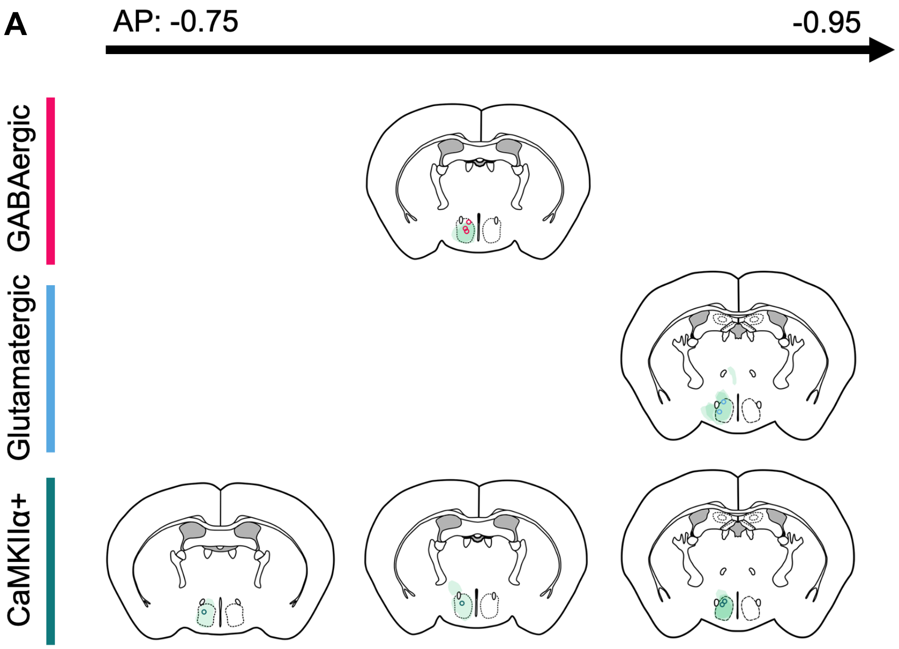


**Figure S1. Viral expression and locations of fibre optic cannulae in the AHN for fibre photometry recordings**

(A) Viral expression depicted by green blots on coronal sections of the mouse brain atlas. Fibre optic cannulae implantation sites indicated by circles.

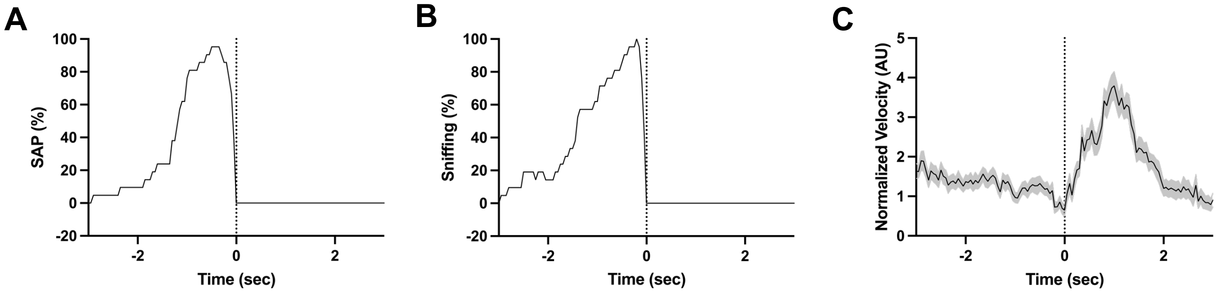


**Figure S2. AHN GABAergic, glutamatergic, and CaMKIIa+ neurons are activated during innate defensive behaviors**

(A) Percent animals demonstrating stretch attenuated postures (SAP) during approach and escape bouts (n = 21)

(B) Percent animals demonstrating sniffing behaviors during approach and escape bouts (n = 21

(C) Normalized velocity during approach and escape bouts (n = 36)

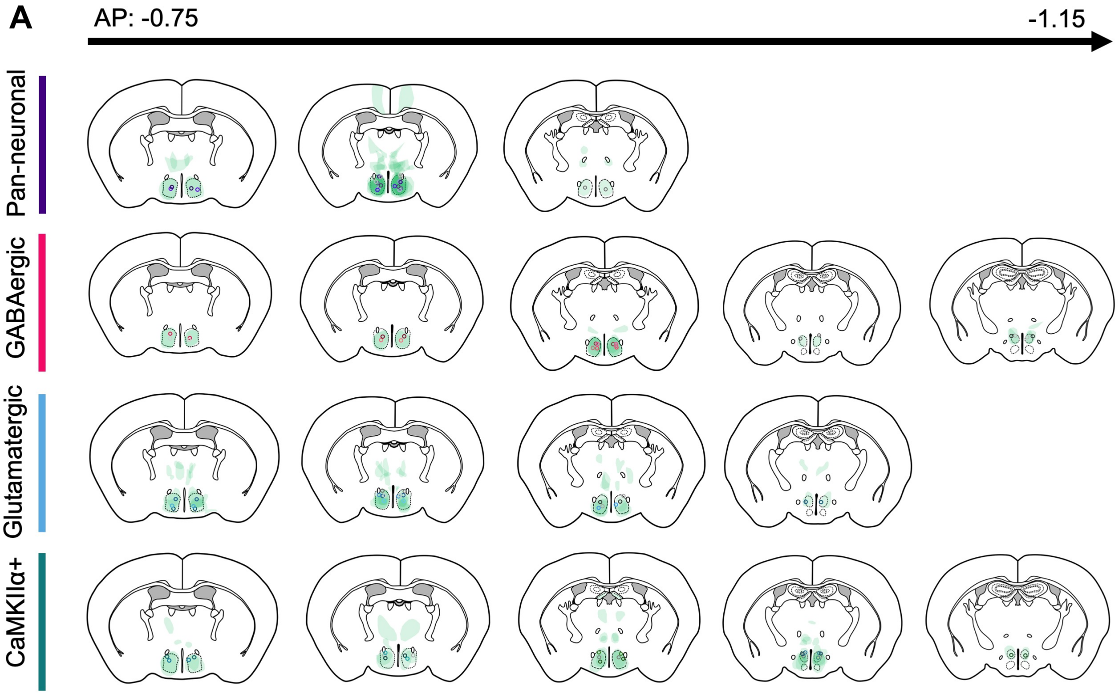


**Figure S3. Viral expression and locations of fibre optic cannulae in the AHN for optogenetic studies**
(A) Viral expression depicted by green blots on coronal sections of the mouse brain atlas. Fibre optic cannulae implantation sites indicated by circles.

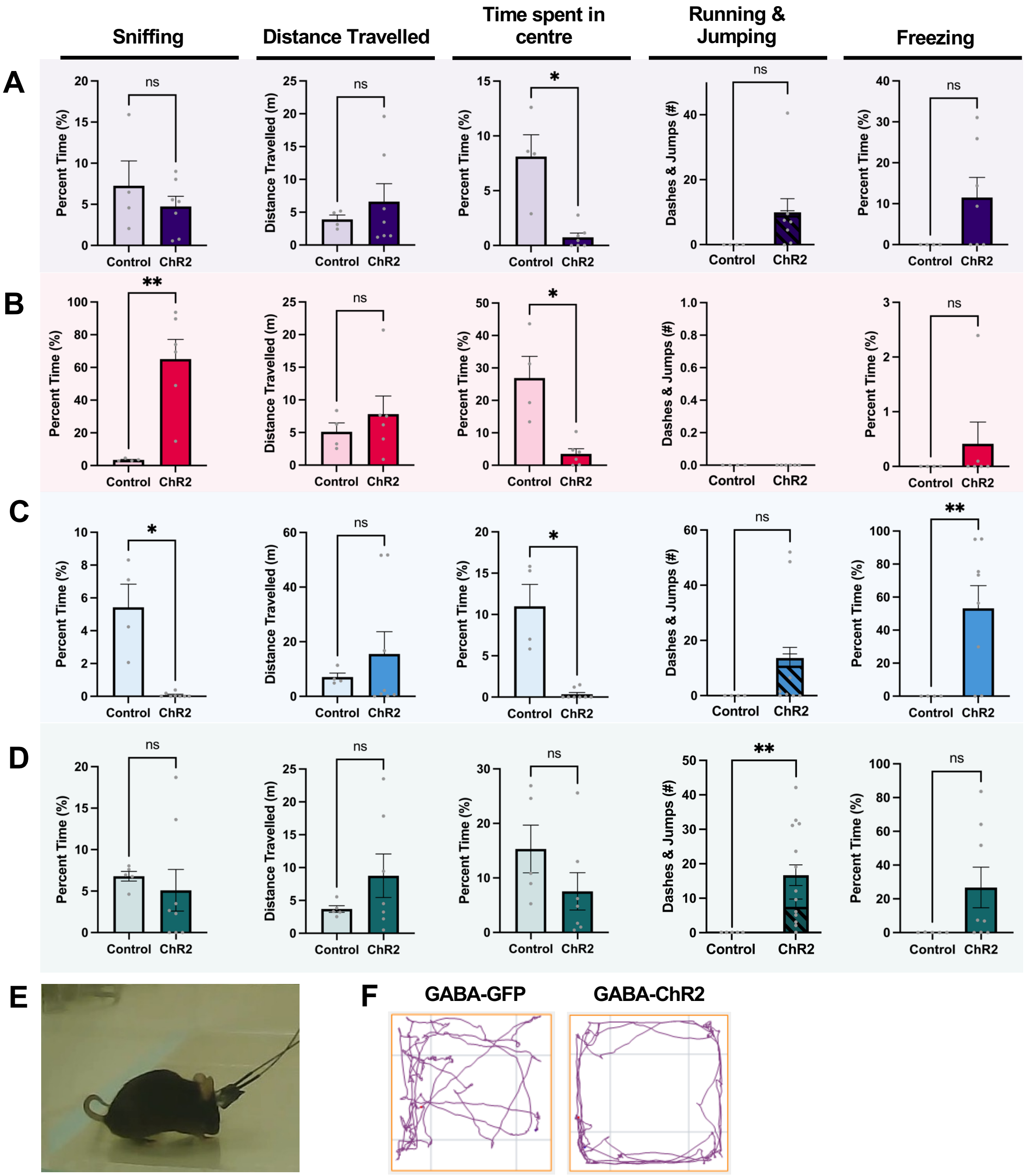
Sniffing
Distance Travelled
Time spent in centre
Running & Jumping
Freezing
A
B
C
D
Percent Time (%)
Distance Travelled (m)
Dashes & Jumps (#)
Control
ChR2
ns
*
**
E
F
GABA-GFP
GABA-ChR2

**Figure S4. High frequency activation of AHN GABAergic, glutamatergic and CaMKIIa+ neurons evoke distinct innate defensive behaviours**

(A) Comparisons of AHN pan-neuronal optogenetic stimulation effects between fluorophore controls ($n = 4$ mice) and ChR2 ($n = 7$ mice) mice on (from left to right) sniffing (Welch's t-test $p > 0.05$), distance travelled (Welch's t-test $p > 0.05$), time spent in centre (Welch's t-test $^*p < 0.05$), running and jumping (Welch's t-test $p > 0.05$), and freezing (Welch's t-test $p > 0.05$) responses during the light ON epoch.

(B) Comparisons of AHN GABAergic neuronal optogenetic stimulation effects between fluorophore controls ($n = 4$ mice) and ChR2 ($n = 7$ mice) mice on (from left to right) sniffing (Welch's t-test $^*p < 0.01$), distance travelled (Welch's t-test $p > 0.05$), time spent in centre (Welch's t-test $^*p < 0.05$), running and jumping, and freezing (Welch's t-test $p > 0.05$) responses during the light ON epoch.

(C) Comparisons of AHN glutamatergic neuronal optogenetic stimulation effects between fluorophore controls ($n = 4$ mice) and ChR2 ($n = 8$ mice) mice on (from left to right) sniffing (Welch's t-test $^*p < 0.05$), distance travelled (Welch's t-test $p > 0.05$), time spent in centre (Welch's t-test $p > 0.05$), running and jumping (Welch's t-test $p > 0.05$), and freezing (Welch's t-test $^{**}p < 0.01$) responses during the light ON epoch.

(D) Comparisons of AHN CaMKIIa+ neuronal optogenetic stimulation effects between fluorophore controls ($n = 5$ mice) and ChR2 ($n = 13$ mice) mice on (from left to right) sniffing (Welch's t-test $p > 0.05$), distance travelled (Welch's t-test $p > 0.05$), time spent in centre (Welch's t-test $p > 0.05$), running and jumping (Welch's t-test $^{**}p < 0.01$), and freezing (Welch's t-test $p > 0.05$) responses during the light ON epoch.

(E) Representative image of a mouse in sniffing posture during GABAergic-ChR2 optogenetic simulation

(F) Representative top-down trajectory plots during a light ON stimulation session (20 Hz) for GABAergic-GFP and GABAergic-ChR2 mice

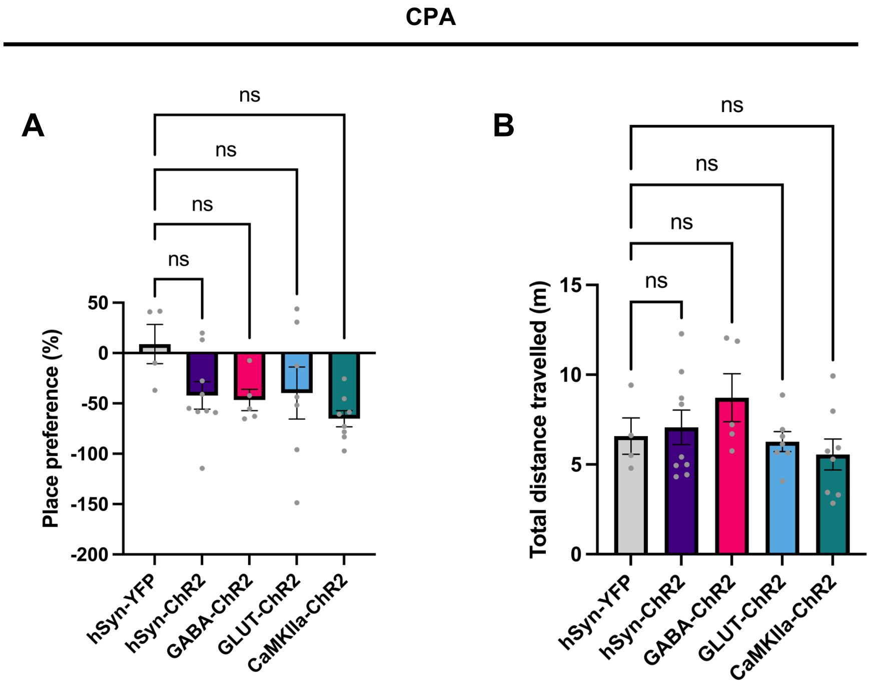


**Figure S5. Activation of AHN GABAergic, glutamatergic, and CaMKIIa+ neurons induces mild conditioned place aversion**

(A) Percent place preference for the stimulation chambers during the CPA sessions by hSyn-GFP (n = 4 mice), hSyn-ChR2 (n = 9 mice), GABA-ChR2 (n = 5 mice), GLUT-ChR2 (n = 7 mice), and CaMKIIa-ChR2 (n = 8 mice) mice (One-way ANOVA $F(4, 28) = 2.009$, $P > 0.05$, with Tukey's multiple comparisons test, $p > 0.05$)

(B) Total distance travelled during the CPA sessions by hSyn-GFP (n = 4 mice), hSyn-ChR2 (n = 8 mice), GABA-ChR2 (n = 5 mice), GLUT-ChR2 (n = 7 mice), and CaMKIIa-ChR2 (n = 8 mice) mice (One-way ANOVA $F(4, 27) = 1.328$, $P > 0.05$, with Tukey's multiple comparisons test, $p > 0.05$)
